# Engineering human myelin microphysiological systems for testing patient treatment response in multiple sclerosis

**DOI:** 10.64898/2026.09.12.750449

**Authors:** Chunhui Tian, Zheng Ao, Hongwei Cai, Jiansen Wang, Nian Wang, Jason Tchieu, Hui-Chen Lu, Ken Mackie, Mingxia Gu, Feng Guo

## Abstract

Multiple sclerosis (MS) is an autoimmune disease of the central nervous system characterized by neuroinflammation, demyelination, and neurodegeneration, associated with a complex interplay between the innate and adaptive immune systems. Currently, no cure is available for MS, and personalized disease-modifying treatment remains largely limited, partially due to the lack of preclinical human models that can faithfully recapitulate disease pathology and evaluate treatment responses at the individual-patient level. Here, we report a human myelin microphysiological system (myelin MPS) platform that recaptures disease phenotype and treatment response for testing patient treatment response. By culturing neural organoids on 3D-printed devices containing directional microfibers, followed by coculture with oligodendrocyte progenitor cells, 96 myelin MPS models can be generated within a conventional well plate. Using this myelin MPS platform, autologous T cells and monocytes from MS patients induced substantially greater demyelination than those from healthy donors, accompanied by expansion of proinflammatory T-cell subsets and increased myelin uptake by monocytes/macrophages after coculture with these healthy myelinating neural tissues. Integration of imaging and flow-cytometric features distinguished healthy-donor, untreated-MS, responder, and nonresponder profiles following treatment with prednisone, glatiramer acetate, interferon β-1a, or dimethyl fumarate. Thus, the myelin MPS platform provides a scalable, human pathophysiology-relevant platform for functional phenotyping and individualized treatment-response evaluation in MS.

## Introduction

Multiple sclerosis (MS) is a chronic autoimmune disease of the central nervous system (CNS) in which persistent inflammation drives demyelination and may ultimately lead to neurodegeneration^1,2^. Demyelination disrupts neural conduction and represents a defining pathological feature of MS. Despite extensive studies, the mechanisms initiating and sustaining demyelination remain incompletely understood. Proinflammatory myelin-reactive T cells and myelin-laden macrophages are closely associated with active demyelination, suggesting that interactions between adaptive and innate immune cells play important roles in myelin injury^3-5^. Currently, no cure is available for MS. Current disease modifying treatments for MS include corticosteroids, such as high-dose oral prednisone for acute inflammatory control and disease-modifying therapies (DMT) such as glatiramer acetate (GA), interferon β-1a (IFNβ-1a), and dimethyl fumarate (DMF)^6,7^. However, therapeutic efficacy and adverse effects differ among patients^8,9^. Such variability can lead to repeated treatment switching and irreversible neural damage^10-12^. Thus, preclinical human models that can faithfully recapitulate disease pathology and evaluate treatment responses are urgently needed but remain lacking.

Human neural organoids are three-dimensional (3D), stem cell-derived tissues that recapitulate key cellular and structural features of the human brain and are increasingly recognized as human-relevant New Approach Methodologies (NAMs)^13-16^. Compared with conventional 2D cultures, their 3D architecture and diverse neural cell populations, better reproduce complex cell–cell interactions and key features of the brain microenvironment^15,17^. Compared with animal models, their human cellular origin preserves human-specific molecular and cellular characteristics, thereby reducing species-related translational barriers^18^. Meanwhile, patient-derived components can be incorporated to investigate patient-specific disease mechanisms and therapeutic responses^19,20^. Previous organoid-based MS models have mainly examined patient-intrinsic neural abnormalities, soluble cerebrospinal fluid–mediated neuroinflammation, or chemically induced demyelination^21-23^. However, models directly integrating living patient-derived T cells and monocytes with myelinated human neural tissue remain limited. In addition, myelin within conventional myelinated organoids is often heterogeneous and difficult to visualize, while organoid variability and low throughput restrict standardized drug screening.

Here, we developed a myelin microphysiological system (myelin MPS) platform to model patient immune cell–mediated demyelination and evaluate personalized treatment responses in MS. By incorporating T cells and monocytes from patients with MS, we reproduced patient-specific immune–myelin interactions and evaluated responses to prednisone, GA, IFNβ-1a, and DMF. Our model has three major advantages. First, two neural organoids are positioned on opposite sides of a 3D-printed device, while electrospinning fibers on the bottom guide the formation of aligned and easily observed myelinated axonal tracts. Second, placing two intact neural organoids ≤ 500 μm apart creates a semi-3D microenvironment that supports close tissue–tissue interactions, thereby more closely recapitulating the spatial organization and multicellular complexity of the human brain. Third, the 3D-printed scaffolds improve standardization and reproducibility of the system, while compatibility with commonly used 96-well plates supports high throughput drug screening. Thus, the myelin MPS establishes a scalable, human-relevant platform for individualized therapeutic evaluation, advancing personalized medicine for MS and other neurological diseases involving myelin dysfunction.

## Results

### Engineer a myelin MPS platform for testing MS treatment response

Multiple sclerosis (MS) is considered an autoimmune disease related to auto-reactive T cell activation, CNS infiltration, and their activation of local myeloid cells, including infiltrating monocytes, which mediate demyelination. Myelin-laden macrophages (MΦ) are key pathological features associated with demyelination in MS^24^. It was described that macrophage and other antigen-presenting cells (APC) could present auto-antigens such as myelin basic protein (MBP) to autoreactive T cells and promote activation^25,26^. In response, T cells will secrete pro-inflammatory cytokines, such as interferon-gamma (IFN-γ), to induce a phagocytic program from macrophages that target the removal of myelin from axons^27,28^. To recapitulate this process *ex vivo*, we develop a patient-specific myelin microphysiological system (myelin MPS) for functional evaluation of MS therapies. T cells and monocytes isolated from MS patients are introduced into human myelinated neural tissue to reproduce patient immune cell–induced demyelination. Clinically relevant MS drugs are then tested in parallel. Effective treatments are identified by their ability to prevent or reduce myelin damage, whereas ineffective treatments fail to prevent demyelination. The resulting personalized drug response report can help prioritize potentially effective therapies and advance personalized treatment for MS (**Fig. 1a**). The myelin MPS consists of two human neural organoids positioned within a 3D-printed device and interconnected by aligned myelinated neural tracts (**Fig. S1**). The ≤500-μm distance between the two human cortical organoids creates a semi-3D brain microenvironment that better recapitulates the spatial organization and multicellular interactions of human brain tissue. The precisely defined device geometry standardizes organoid placement and inter-organoid distance, thereby improving experimental consistency and reproducibility. In addition, its compatibility with conventional 96-well plates enables parallel culture, drug treatment, and analysis, supporting high-throughput drug screening (**Fig. 1b & 1c**). Briefly, 3D human embryonic stem cell (hESC) derived neural organoids were seeded onto uniform electrospinning nanofibers with 2-4 μm diameters. Once seeded on-chip, neurons from neural organoids will migrate onto nanofibers to cover them in 2 weeks. Upon successful migration, we will subsequently introduce hESC-derived oligodendrocyte progenitor cells (OPCs) on our chip to provide a substrate for myelination that can be achieved in an additional 2 weeks. Myelination of the neurons could be visualized by red FluoroMyelin dye (**Fig. 1d**). The semi-3D brain microenvironment provided by organoids facilitates the myelination to generate more uniform myelination with higher myelin coverage as compared with OPCs alone on nanofibers (**Fig. S2**). The myelinated neuronal cultures were then kept on chip before the seeding of the immune cells.

**Fig 1.**
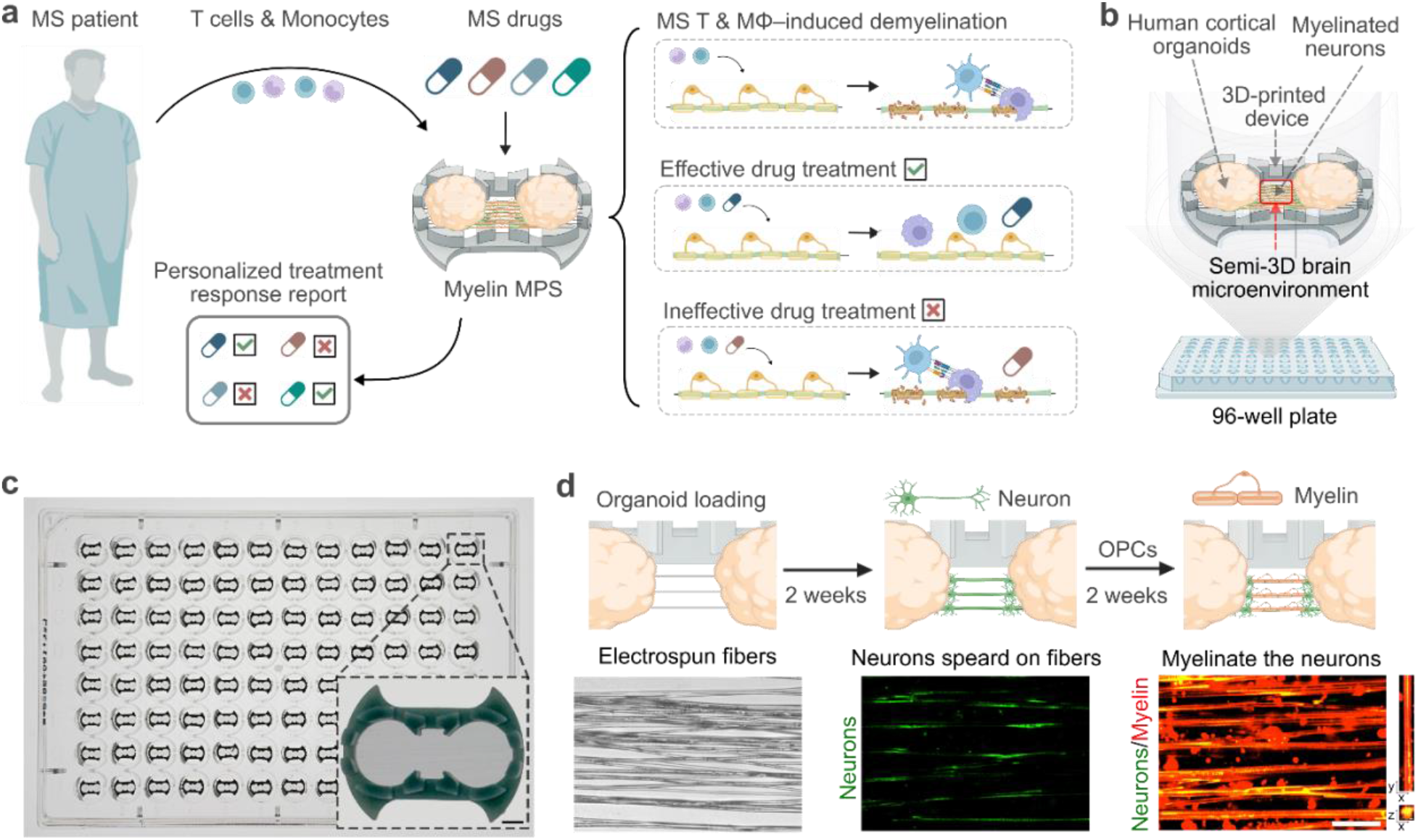
Myelin MPS platform for testing MS treatment response. (**a**) T cells and monocytes isolated from MS patients are introduced into the myelin MPS allowing observation of on-chip demyelination and tested with different MS treatments to identify effective and ineffective therapies. (**b**) A single myelin MPS consists of two cortical organoids positioned on opposite sides of a 3D-printed scaffold, which supports the growth of myelinated neurons and creates a semi-3D brain microenvironment. (**c**) Image of the myelin devices in a commonly used 96-well plate and an enlarged view of the device. Scale bar: 500 μm. (**d**) Formation of the myelinated neural network within the device. Human cortical organoids positioned on opposite sides of the scaffold extend aligned neurites through the central channels, followed by myelin formation along the neurites. Representative bright-field and fluorescence images show aligned neurites (green) and myelin (red). Scale bar: 50 μm.

### Characterize myelin MPS for modeling demyelination

To test whether we could observe on-chip demyelination mediated by T-monocyte interactions on-chip, we tested our model with MBP reactive T cells and THP-1 monocytic cells as positive controls. MBP reactive T cells were purchased from a commercial vendor, which were expanded from an MS patient with a significant risk human leukocyte antigen (HLA) haplotype: DRB1*15:01, using the matching B lymphoblastoid cell lines (B-LCL) and MBP antigen. To match the HLA haplotype, we chose the THP-1 cell line, which is a human monocytic cell line with the HLA-DRB1*15:01 haplotype^29^. Generally, differentiation of THP-1 towards macrophage requires prolonged adherent culture with stimulants such as phorbol 12-myristate 13-acetate (PMA)^30^. Here we found that, unlike 2D cultures, the semi-3D neuronal culture environment is sufficient to induce THP-1 into a macrophage phenotype as indicated by increased CD68 expression as well as extended morphology on-chip (**Fig. 2a**). Adhered THP-1 in our system could further respond to pro-inflammatory signaling such as IFN-γ treatment on-chip as indicated by increased CD68 expression on THP-1 cells (**Fig. 2b**). Furthermore, IFN-γ treatment was able to induce phagocytosis of myelin as indicated by intracellular MBP staining in activated THP-1 cells (**Fig. 2c**). We then loaded the chip with THP-1 and MBP reactive T cells to test the T-MΦ interaction and their ability to perform demyelination on our device. We found that once MBP-reactive T cells were co-loaded with THP-1 cells, they could induce significant demyelination on-chip, as indicated by live imaging of demyelination via FluoroMyelin dye labeling (red) (**Fig. 2d**). Quantification data also indicated that MBP-reactive T cells could induce demyelination to a greater extent as compared with IFN-γ treatment alone (**Fig. 2e**).

**Fig 2.**
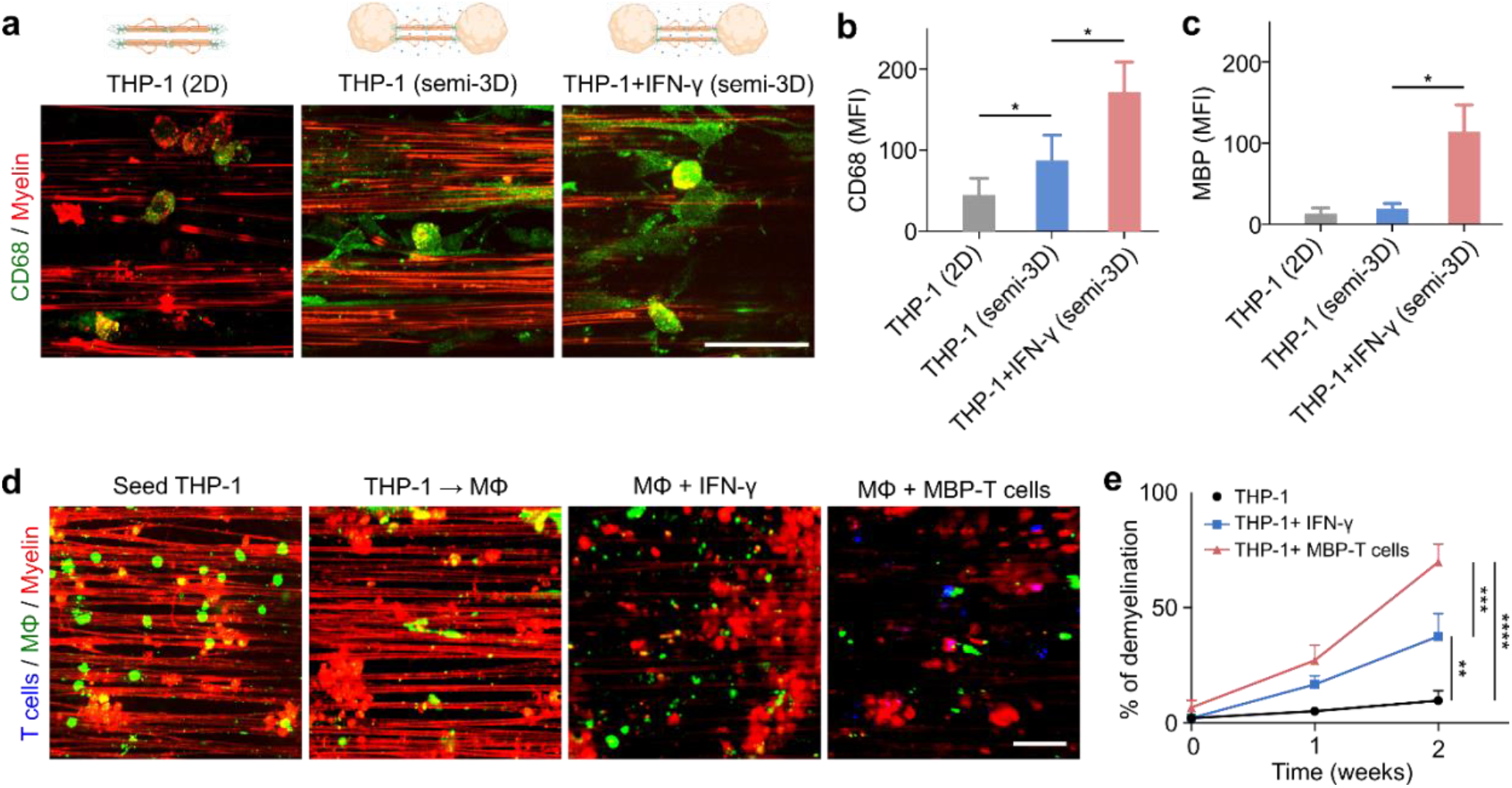
Characterization of the myelin MPS platform for modeling immune cell–mediated demyelination. (**a**) THP-1 seeding and spontaneous differentiation into macrophage morphology in semi-3D myelin MPSs. Post-IFN-gamma induced activation, THP-1 cells rapidly remove myelin from neuron cells. (**b**) Quantification of CD68 expression on THP-1 in 2D, semi-3D and IFN-gamma activated (semi-3D) culture conditions after 24 hours coculture. (**c**) Quantification of intracellular MBP staining withing THP-1 cells. (**d**) Immunofluorescence staining of CD68, a M1 macrophage (MΦ) marker to confirm THP1 differentiation and polarization. (**e**) Quantification of demyelination in THP-1, THP-1 + IFN-gamma and THP-1+MBP-T cells conditions in semi-3D myelin MPSs. Mean ± SEM, *n* = 3 samples, from three independent experiments. Scale bar: 50 μm.

### Characterize demyelination by T cell and monocytes from MS patients

Current functional phenotyping assays of PBMCs such as ELISPOT rely on interactions between antigen-presenting monocytes and antigen-reactive T cells to stimulate T cells’ production of functional cytokines, such as IFN-γ, to detect antigen-reactive T cell frequencies^31^. However, frequencies of such antigen-reactive T cells do not always correlate with disease onset and severity due to the low frequency of autoreactive T cells in peripheral blood and is compounded by the multifaceted and dynamic changes of T cell functions modulated by the local brain microenvironment and immune interactions^32^. *In vivo*, MS disease pathology consists of a cascade of events including autoreactive T cell infiltration into CNS, recruitment of monocytes to demyelination edge, and activation of monocytes/macrophages, which mediate myelin phagocytosis and removal^33^. Single readout assays such as IFN-γ secretion from T cells do not reflect the complex events at play. Thus, we sought to test whether our myelination MPS could be used to interrogate such a dynamic demyelination process. We loaded MS and healthy donors’ PBMC-derived T cells with autologous monocytes onto our myelin MPS to observe their interactions. As controls, we loaded chips with T cells or monocytes as single populations. We found that MS T cells alone could only induce moderate demyelination on-chip (demyelination percentage: 14.7% ± 3.4%), whereas MS monocytes alone did not induce significant demyelination compared to controls. However, when MS T cells and monocytes were added together on-chip, significant demyelination of 67.6% ± 19.7% could be observed (**Fig. 3a**). In contrast, healthy donor-derived T and monocytes co-culture (as an *in vitro* model on-chip) induced a much lower extent of demyelination 27.1 ± 12.4% (n=5 for all groups) (**Fig. 3b**). We next asked whether immune cell density was associated with sites of demyelination on-chip. We quantified T cell and monocyte numbers as well as their co-localization in the *in vitro* model and observed proliferation/persistence of T cells in MS group as indicated by increased T cell number after 2 weeks of co-culture. In contrast, monocytes numbers did not deviate from initial seeding. Moreover, we found a significantly higher spatial colocalization of T cell and monocytes in the MS group as compared with HD group indicating potential T-monocyte interactions (**Fig. 3c**). Finally, we interrogated by flow cytometry, the T cell and monocytes subpopulations. We found that IFN-γ+ Th1 CD4 T cells and IL-17+ Th17 CD4 T cells were selectively amplified during on-chip incubation in the MS group, as well as TNF-α+ CD8 T cells group. Regarding CD14+ myeloid cells, we analyzed intracellular MBP staining to indicate macrophage phagocytosis of myelin. Indeed, MS patients’ monocytes have significantly higher intracellular MBP staining post-on-chip incubation as compared to those from healthy donors (**Fig. S3**).

**Fig 3.**
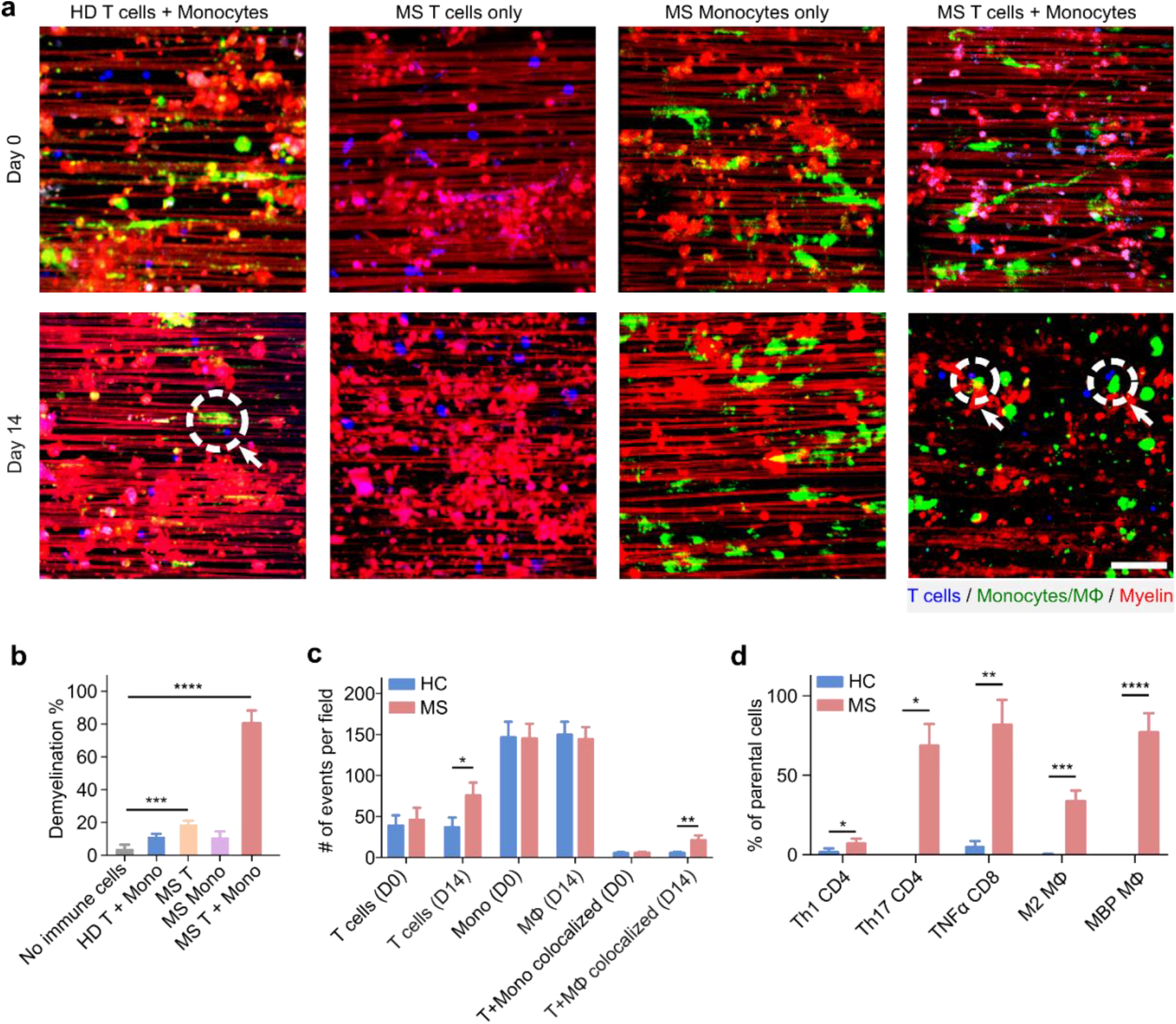
Characterization of demyelination by T cells and monocytes from MS patients. (**a**) Day 0 and Day 14 images of semi-3D myelin MPSs seeded with HD T + monocytes, MS T cells, MS monocytes, and MS T + monocytes conditions. (**b**) Quantification of Day 14 demyelination percentages under no immune cells control, HD T cells + monocytes (HD T + Mono), MS T cells (MS T), MS monocytes (MS Mono), and MS T cells + monocytes (MS T + Mono). (**c**) Quantification of T cells number, monocyte number and co-localization of T and monocytes per field of view from HD and MS cells seeded devices. (**d**) Flow cytometry quantification of different immune cell compartments in HD and MS devices 14 days post immune cell seeding. Mean ± SEM, *n* = 3 samples, from three independent experiments. Scale bar: 50 μm.

Additionally, we investigated the activation state of the monocytes by measuring CD40 versus CD163 expression, as previous reports from human MS studies indicated these are the two consistent markers for M1 and M2 states in monocytes/macrophages^34^. We found that after co-incubation on-chip, monocytes from MS patients were polarized towards M2 (CD163+ CD40-) (**Fig. S4**). This is consistent with previous reports that phagocytosis of myelin will polarize monocytes towards M2 phenotypes (**Fig. 3d**)^35^. Overall, we found that on-chip co-incubation of MS peripheral blood-derived T cell and monocytes is sufficient to induce demyelination, which is a distinctive functional phenotype not observed from healthy donors’ immune cells. Additionally, our control experiments have revealed that the on-chip demyelination is amplified by T cell-monocyte interaction, removal of 1 component from this pair could significantly reduce the demyelination on-chip.

### Evaluate patient-specific treatment response in MS

Current treatments in MS rely on immune-modulatory drugs that suppress the immune system. Selecting the right initial disease-modifying therapy (DMT) for MS could greatly benefit disease outcomes^12^. However, the mechanisms of action for most MS treatments remain largely elusive, making it difficult to select the right treatments for each patient to maximize efficacy early on^36^. Prior studies have shown that genetic makeup and phenotypes of peripheral immune cells could be utilized to inform personalized MS therapy^37^. Hence, we tested 4 standard-of-care MS treatments on-chip to observe their effects on immune cell phenotypes and their function to slow down the demyelination process from 6 DMT treatment naive patient samples (**Fig. 4a**). To evaluate on-chip phenotypes of immune cells post-on-chip incubation, we profiled the immune cells by flow cytometry. We characterized CD4+ T cells by surface markers into naïve CD4 (CD45RA+ CD45 RO-), memory CD4 (CD45RA-CD45RO+), effector memory CD4 (CD45RA-CD45RO+ CCR7-), and similarly for CD8 T cells. For monocytes (CD14+), we further gated these cells as M1 (CD40+ CD163-) or M2 (CD40-CD163+) states. For intracellular staining, we profiled CD4+ T cells into IFN-γ+ Th1, IL4+ Th2, and IL-17+ Th17. For CD8 T cells, we profiled them into IFN-γ+ CD8 and TNF-α+ CD8. Additionally, we determined the levels of intracellular staining of MBP+ to measure levels of phagocytosis in monocytes/macrophages (**Fig. S5**). Combining these 14 parameters from flow cytometry (**Table S3**), we utilized unsupervised dimension reduction algorithm -Uniform Manifold Approximation and Projection (UMAP) to project the flow cytometry analysis data from 5 healthy control samples and 6 MS samples (untreated conditions) onto a x-y 2D plane.

**Fig 4.**
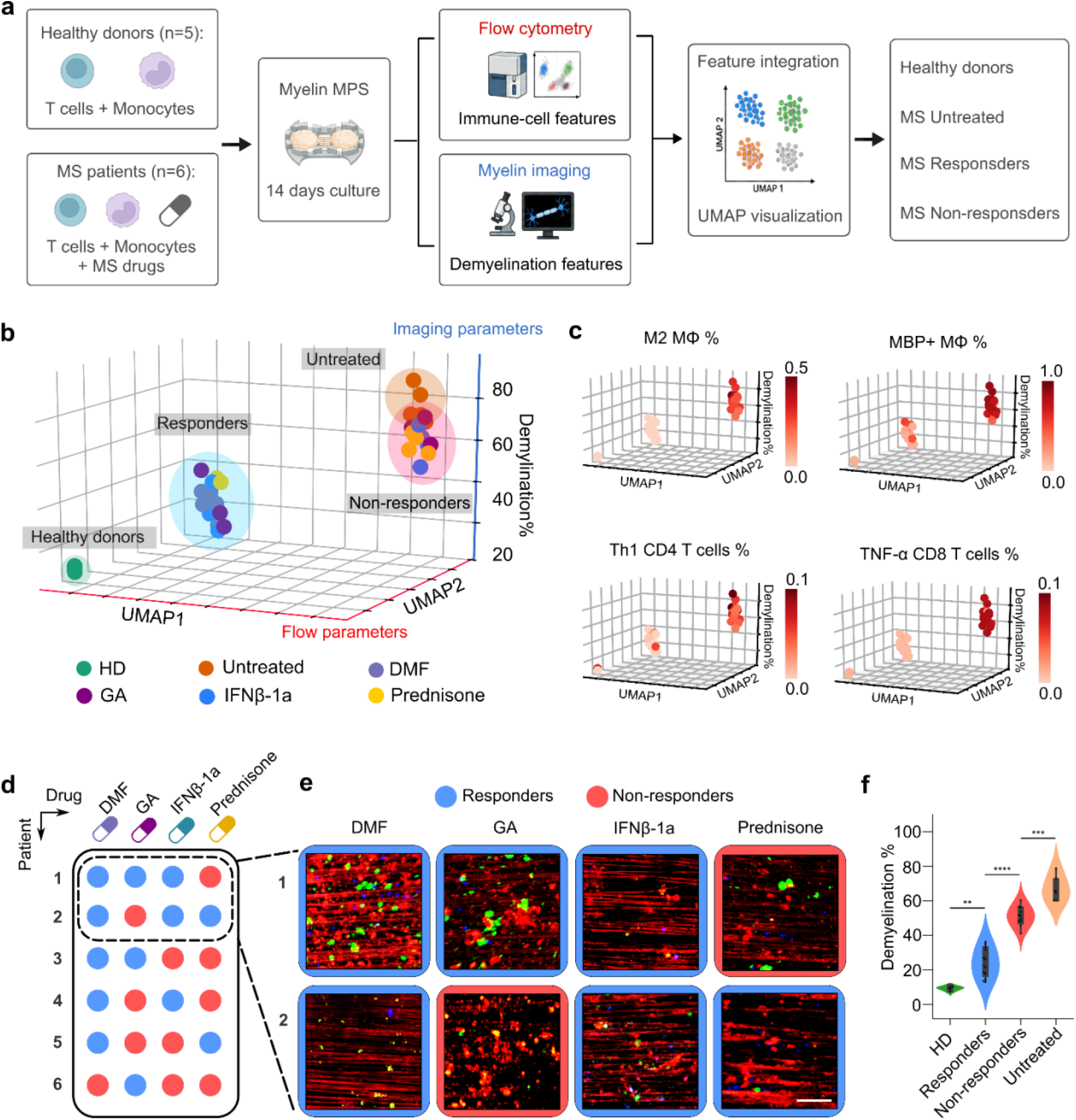
Evaluation of patient-specific treatment response in MS. (**a**) A schematic showing the experiment flow of evaluating patient-specific treatment response in MS. (**b**) 3D visualization healthy donor (HD), untreated and treated functional phenotypes of immune cells derived from PBMC on-chip. Flow cytometry derived parameters were projected onto x and y axis by UMAP, and imaging-based demyelination quantification was visualized on z-axis. (**c**) Visualization of different cell population compositions projected onto 3D UMAP guided clustering as described in 4b. (**d**) Responses to 4 standard-of-care MS drugs in all 6 MS patients (i to iv). (**e**) Visualization of demyelination in first and second patients with 4 treatments. (**f**) Quantification of demyelination percentages in healthy donor (HD), responders, non-responders, and untreated groups. Scale bar: 50 μm.

The results show that the flow parameters formed 2 distinct clusters on the UMAP projections. By further implementing on a third axis (z-axis) with imaging-based on-chip demyelination data on top of the 2D UMAP plot, we found that the UMAP clusters of healthy donors were distributed within the low demyelination percentages (low on Z-axis), whereas the untreated MS samples were distributed within the high demyelination percentages (high on Z-axis) (**Fig. 4b**). We further tested 4 FDA-approved, first line therapies of MS, including prednisone, glatiramer acetate (GA), interferon β-1a (IFNβ-1a) and dimethyl fumarate (DMF). We projected the immune cell phenotypes post-treatment onto the 3D space with UMAP projection on x-y and demyelination percentage on z. We found that the post-treatment immune cell phenotypes clustered into 2 separate clusters. 1 cluster has lower demyelination on-chip, with UMAP projection closer to healthy donor controls, indicating the treatment effectively reduced on-chip demyelination and promoted an anti-inflammatory phenotype, thus we termed these as responders. On the contrary, the other cluster has higher on-chip demyelination, and clustered closer to the untreated group on the UMAP projection based on flow cytometry parameters, which we named as non-responders. Upon closer examination, we found that the clusters of non-responders and untreated group also have higher MBP+ monocytes with M2 phenotypes, with higher IFN-γ+ Th1 CD4 T cells and TNF-α+ CD8 T cells (**Fig. 4c**). We further confirmed that the on-chip demyelination by imaging (**Fig. 4d & 4e**). We found that the demyelination percentage was the highest in the untreated group, followed by non-responder group. Where this demyelination is much lower in the responder group and lowest with healthy donor samples with statistical difference (**Fig. 4f**).

## Discussion

Traditionally, MS is diagnosed and monitored through costly brain imaging such as magnetic resonance imaging (MRI), or invasive sampling and profiling of cerebrospinal fluid (CSF) components^38,39^. Such testing is usually limited to patients with clear symptoms and as limited utility to be used as companion diagnosis tools which enables repeated, minimally invasive sampling. Additionally, there is a great need to develop liquid biopsy tests to guide transition of first line to second line therapies in MS^40,41^. To make the MS testing more accessible and less invasive, it has always been of great interest to develop peripheral blood-based tests for MS^42-44^. However, sampling and profiling peripheral immune cells for MS disease diagnosis and disease monitoring have been challenging due to the low frequency of autoreactive immune cells in peripheral blood and the confounding low fraction of autoreactive cells also present in healthy donors’ blood. Attempts have been made to distinguish MS blood immune cells from those from healthy donors via detailed compartment analysis via flow cytometry, single cell analysis, or isolation and culture of single clones of cells^45-48^. However, such reports are either inconclusive, sometimes contradicting, or based on case studies that cannot easily be scaled for clinical applications.

Additionally, traditional functional phenotyping analyses of MS peripheral immune cells were often based on *ex vivo* culture experiments with artificial external stimulants such as lipopolysaccharides (LPS) or phorbol 12-myristate 13-acetate (PMA), which were not present in the MS brain^49-51^. To better recapitulate the diseased phenotype of MS immune cells in patients’ brain, here we established a scalable semi-3D myelin MPS to condition the immune cells to better reflect their diseased phenotypes in brain. Studies have indicated that immune cells were constantly reprogrammed by its residing niche and show tissue-specific phenotype^52-54^. Here, we demonstrated that by co-culturing monocytes on our 3D myelination MPS, they can be readily conditioned as macrophages and respond to external stimuli. Additionally, co-addition of T cells and monocytes from MS peripheral blood have recapitulated the key features of demyelination in MS lesions, a distinct phenotype not seen with healthy donors’ blood derived counterparts. Thus, this test holds potential as a diagnostic test for MS. Additionally, different MS therapeutics have distinct mechanism of actions (MoA) which are generally challenging to recapitulate *in vitro*. By integrating components from the local brain microenvironment, innate immune components (monocytes) as well as adaptive immune components (T cells), we successfully demonstrated therapeutic effects of 4 different DMT with distinct MoA^55-57^. This highlighted the sensitivity of our model as well as its utility as a potential drug screening platform.

One disadvantage is the time-consuming steps lead to establishment of the model. Establishing the cortical organoids generally takes 6-8 weeks to allow proper neuron and glia development. Further on-chip culture and myelination steps take another 4 weeks, leading to requirement of preparation time of 10-12 weeks before we could seed immune cells on-chip. Once the model is ready, it could be maintained for up to 2 months in culture. Thus, our current solution is to have staggered timed models for larger scale experiments. Alternatively, we could evaluate if the organoid or OPC mediated myelination in our model could be replaced with simpler models such as neuronal cell line cultures with myelin expression. Another drawback of the model is the lack of other immune cells such as brain residential microglia and B cells, which also play essential roles in MS disease progression and drug responses^58-60^. Additionally, due to the limited information we have on the MS patients. We could not evaluate if the on-chip responses of immune cells to MS disease modifying drugs correlate to the actual patients’ clinical responses. A larger scale clinical trial with carefully designed sample collection before and during various timepoints of MS treatments could help us better answer this question.

Lastly, here we introduced MS patients derived immune cells on-chip. One can envision a different approach where we introduce patient induced pluripotent stem cell (iPSC) derived cortical organoids together with iPSC derived oligodendrocytes. This could render us a model to study the effect of genetic mutations of MS patients on oligodendrocyte migration, survival and myelination capacity under normal and/or stressed conditions^61-63^. To conclude, here we reported a functional phenotyping test using MS peripheral blood immune cells via *ex vivo* 3D culture on-chip. This test holds potential to be widely adapted for disease modeling, disease diagnostics as well as personalized therapy for MS and other demyelinating diseases.

## Materials and Methods

### Culture of THP-1 cells

THP-1 cells were purchased from American Type Culture Collection (ATCC). Cells were maintained in RPMI-1640 Medium supplemented with 10% fetal bovine serum (Gibco) and 0.05 mM 2-mercaptoethanol (Sigma) in a 37°C, 5% CO_2_ humidified incubator. Cells were enumerated every other day and passaged below or at 1 million cells per milliliter, at a ratio of 1:3 into new flasks.

### Culture of MBP-reactive T cells

Anti-myelin basic protein (MBP) T cells were purchased from Cellero (donor #3). MBP-reactive T cells were maintained in RPMI-1640 Medium supplemented with 10% fetal bovine serum (Gibco), 0.05 mM 2-mercaptoethanol (Sigma), 30 IU/mL human recombinant IL-2 (R&D systems) and 1:1 CD3/CD28 Dynabeads (Gibco) in a 37°C, 5% CO_2_ humidified incubator till the day of use.

### Isolation of primary T cells and monocytes from patient or healthy donors’ PBMC

Frozen peripheral blood mononuclear cells (PBMC) from healthy donors or multiple sclerosis patients were purchased from Accelerated Cure Project (ACP) repository. Frozen PBMC aliquots were thawed and resuspended in prewarmed RPMI-1640 Medium supplemented with 10% fetal bovine serum (FBS) (Gibco) and spin down at 300g for 10 minutes. Pelleted cells were resuspended in 1X Phosphate Buffered Saline (PBS) supplemented with 10% fetal bovine serum (FBS). Single cell suspensions were enriched by human pan T cell isolation kit (Miltenyi) for T cells or classical monocyte isolation kit for classical monocytes (Miltenyi).

### Labeling of immune cells

Isolated T cells or monocytes were resuspended in RPMI-1640 medium at 1 million cells per 1 mL to stain with blue CMAC cell tracker (T cells) (with 10% FBS) or Vybrant DiO (monocytes) (Invitrogen) (Serum-free medium) at 1:1,000 dilutions at 37 °C for 1 hour. Labeled cells were then washed twice with RPMI-1640 medium supplemented with 10% FBS before usage.

### Culture of oligodendrocytes

Human embryonic stem cell (WA09) derived oligodendrocyte progenitor cells (OPC) were purchased from Millipore sigma. The OPCs were plated onto 1% growth factor reduce (GFR) Matrigel (Corning) pre-coated T25 tissue flasks in human OPC expansion media (Millipore Sigma). The OPCs were passaged at 70-80% confluency using accutase (Stemcell Technologies).

### Culture of cortical organoids

Human cortical organoids were established as previously described^15,64,65^.Human embryonic stem cells (WA09) were obtained from WiCell under material transfer agreement approved by both WiCell institute and Indiana University. To establish human cortical organoids, 9,000 human embryonic stem cells (WA09, WiCell) were aggregated in EB formation medium. The EBs were then subject to dual-SMAD inhibition by Dorsomorphin and SB431542 for neural induction for 10 days. The spheroids were then subject to neural epithelial expansion in FGF-2 containing medium for 7 days and then EGF+FGF-2 containing medium for 7 days. The spheroids were then further differentiated in BDNF, GDNF, ascorbic acid, cAMP and NT3 containing medium for further neuron and glial maturation. Finally, the mature organoids were maintained in BrainPhys imaging optimized medium for additional 1-6 months before addition to our 3D neuronal chip. Detailed medium compositions can be found in **Table S1**.

### Establishment of 3D myelinated neuronal model

To establish 3D myelinated neuronal model, we first coated the electrospun nanofibers with 0.01% poly-L-ornithine solution (Millipore Sigma) overnight, followed by 20 μg/mL fibronectin coating for >4 hours. We then plated mature (>1.5 months old) human cortical organoids onto the coated nanofibers, and culture them in cortical organoid maintenance media supplemented with 20 ng/mL human brain-derived neurotrophic factor (BDNF) (PeproTech). The neurons from mature cortical organoids would outgrow and extend on nanofibers to fully cover them in 2 weeks. Once the nanofibers were fully covered by outgrown neurons from cortical organoids, we plated OPC onto the nanofiber device at a seeding density of 10,000 OPC per device. The medium was switched to glial medium (**Table S1**) for 2 weeks to allow full myelination of the device. To visualize myelination, device was stained with FluoroMyelin red fluorescence myelin stain (Invitrogen) at 1:300 in glial medium for 30 minutes in 37 °C incubator and washed twice.

### Observation of T-monocytes interactions on chip

To observe T and monocytes interactions on-chip, labeled T and monocytes were plated on-chip at 5,000 cells each. Chips were imaged using an inverted Olympus spinning disk microscope to visualize cell-cell interactions and demyelination process.

### Flow cytometry analysis of primary immune cells

At day 14 post-on-chip incubation, immune cells were dissociated from 3D neuronal cultures by accutase treatment at 37 °C for 6 minutes. Digested single-cell suspensions were resuspended in flow buffer made with filtered 1X phosphate-buffered saline (PBS, Gibco), supplemented with 10% fetal bovine serum (FBS, Gibco). The cells were then labeled with corresponding fluorophore-conjugated antibodies for 30 minutes at 4 degrees (**Table S3**). For intracellular cytokine staining, we initially treated immune cells on-chip with BD GolgiStop protein transport inhibitor for 6 hours. The cells were then dissociated from the chip by Accutase (STEMCELL Technologies) and stained using BD Fixation/Permeabilization Kit. Compensation controls were prepared using anti-rat or anti-mouse compensation particles (BD Biosciences). Compensation controls and fluorescent minus one control were run together with the experimental samples using a BD LSRII flow cytometer. Flow cytometry data was analyzed using FlowJo (v10).

### Drug treatment of MS models

For drug treatments, prednisone (Cayman chemicals) was applied at a concentration of 35.8 ng/mL, glatiramer acetate (Cayman chemicals) was applied at a concentration of 20 μg/mL, dimethyl fumarate (Cayman chemicals) was applied at a concentration of 3.6 mg/mL and human recombinant interferon beta 1 alpha (Bio-rad) was applied at a concentration of 10 ng/mL. Drugs were diluted in glial medium and treated for 2 weeks.

### Immunofluorescence staining

For immunofluorescence staining, samples were treated with 4% paraformaldehyde overnight, wash twice with 1X PBS, and treated with 2N HCl for 15 minutes. The treated samples were then washed 3 times with 1XPBS, followed by blocking/permeabilization with blocking buffer (1X PBS + 3% bovine serum albumin + 0.3% triton X-100) for 1 hour. The samples were then incubated with primary antibodies overnight at 4 °C, washed 3 times with 1X PBS, then stained with secondary antibodies at room temperature for 1 hour in dark. The samples were finally washed 3 times with 1X PBS and stained with DAPI and coverslipped with prolong gold anti-fade mounting medium (Invitrogen). The samples were visualized using an inverted Olympus microscope (IX-83) or confocal microscope (Leica SP8).

### Statistical analysis

All data were extracted and analyzed using Prism 7 (GraphPad Software). P-value between 2 samples were analyzed by student’s t-tests. P-value among 3 or more samples were analyzed by one-way ANOVA followed by Tukey’s honestly significant difference (HSD) post hoc test. P-values were denoted as following: * <0.05; ** <0.01; *** <0.005; **** < 0.001.

## Acknowledgment

F.G. acknowledges Indiana University departmental start-up funds, and the National Institute of Health Awards (DP2AI160242, U54AG090792, and R01GM160423). We also wanted to acknowledge Dr. Alline C Campos for her input.

